# The time to peak blood bicarbonate (HCO_3_^−^), pH, and strong ion difference (SID) following sodium bicarbonate (NaHCO_3_) ingestion in highly trained adolescent swimmers

**DOI:** 10.1101/2021.03.01.433339

**Authors:** Josh W. Newbury, Matthew Cole, Adam L. Kelly, Richard J. Chessor, S. Andy Sparks, Lars R. McNaughton, Lewis A. Gough

**Affiliations:** Human Performance and Health Research Group, Centre for Life and Sport Sciences (CLaSS), Department of Sport and Exercise, Birmingham City University, Birmingham, UK; Sports Science and Sports Medicine Team, British Swimming, Loughborough, Leicestershire, UK; Sports Nutrition and Performance Research Group, Department of Sport and Physical Activity, Edge Hill University, Ormskirk, UK

## Abstract

**Background:** Contemporary research suggests that the optimal timing of sodium bicarbonate (NaHCO_3_) should be based upon an individual time in which bicarbonate (HCO_3_^−^) or pH peaks within the blood. However, the mechanisms surrounding acidosis on exercise performance are contested, therefore it is plausible that the ergogenic effects of NaHCO_3_ are instead a result of an increased strong ion difference (SID) following ingestion. Since the post-ingestion time course of the SID is currently unknown, the purpose of this study was to investigate the pharmacokinetics of the SID in direct comparison to HCO_3_^−^ and pH.

**Methods:** Twelve highly trained, adolescent swimmers (age: 15.9 ± 1.0 yrs, body mass: 65.3 ± 9.6 kg) consumed their typical pre-competition nutrition before ingesting 0.3 g·kg BM^-1^ NaHCO_3_ in gelatine capsules. Capillary blood samples were then taken during quiet, seated rest on nine occasions (0, 60, 75, 90, 105, 120, 135, 150, and 165 min post-ingestion) for the assessment of time course changes in HCO_3_^−^, pH, and the SID.

**Results:** On a group mean level, no differences were found in the time in which each variable peaked within the blood (HCO_3_^−^ = 130 ± 35 min, pH = 120 ± 38 min, SID = 96 ± 35 min; p = 0.06). A large effect size was calculated between the timing of peak HCO_3_^−^ and the SID (*g* = 0.91), however, suggesting that a difference may occur between these two measures in practice.

**Conclusions:** A time difference between peak HCO_3_^−^ and the SID presents an interesting avenue for further research since an approach based upon individual increases in extracellular SID has yet to be investigated. Future studies should therefore compare these dosing strategies directly to elucidate whether either one is more ergogenic for exercise performance.

## Introduction

Sodium bicarbonate (NaHCO_3_) is recommended to athletes based upon adequate evidence of a performance enhancing effect [1]. It is well acknowledged that NaHCO_3_ ingestion increases buffering capacity by increasing bicarbonate (HCO_3_^−^) and pH concentrations within the blood, although the magnitude of these increases is variable between individuals [2,3]. Increasing blood alkalosis alters the pH gradient between the intracellular and extracellular compartments, subsequently leading to the upregulation of the lactate–hydrogen ion (H^+^) co-transporter to efflux acidic H^+^ from the active musculature and into circulation [4,5]. During exercise, accelerating the removal of H^+^ is purported to offset fatigue since intracellular acidosis is associated with debilitating effects on the capacity to sustain muscle force production [6–10]. More specifically, cellular acidosis is suggested to reduce muscle shortening velocity and cross-bridge cycling by inhibiting calcium ion (Ca^2+^) sensitivity [6,9] and myosin ATPase activity [10,11], whereas additional impairments of key glycolytic enzymes [12,13] and the strong ion difference (SID) [7,14,15] could reduce the available ATP substrates and action potentials necessary to maintain muscular contractions, respectively. Due to the complexity of fatigue, the role of acidosis in each of these mechanisms is contested as a causative factor [16–18]. However, since NaHCO_3_ continues to provide a strong ergogenic benefit to exercise, further investigation is warranted concerning the biochemical changes that occur following ingestion.

There is an apparent increase in the ergogenic potential of NaHCO_3_ when blood HCO_3_^−^ increases above baseline concentrations by 5 mmol·L^-1^, whereas increases above 6 mmol·L^-1^ are associated with almost certain performance enhancement [19,20]. However, the time taken to reach these thresholds varies considerably when NaHCO_3_ is consumed in either capsule (range: 40 – 240 min [3,21,22]), or solution form (range: 40 – 125 min [2,23–25]), therefore highlighting a potential flaw in the current dosing guidelines (i.e., 60 – 150 min pre-exercise for all athletes [1]). Contemporary research is addressing this issue by timing exercise to coincide with an individual peak in HCO_3_^−^ concentration [21,23–25], though at present, only Boegman et al. [21] has directly compared exercise performance between NaHCO3 ingested at an individual peak in HCO3^−^ (40 – 160 min pre-exercise) versus a traditionally recommended dosing protocol (60 min pre-exercise for all athletes). The individualised approach significantly improved the rowing performance (2,000 m time-trail) of 18 out of 23 world-class rowers (Individualised = 367.0 ± 10.5 s vs. Traditional = 369.0 ± 10.3 s), however, this result occurred with only a small difference in pre-exercise HCO_3_^−^ (Individualised = +6 mmol·L^-1^ vs. Traditional = +5.5 mmol·L^-1^). It is therefore plausible that another mechanism besides those associated with increased HCO_3_^−^ is involved with the ergogenic properties of NaHCO_3_ supplementation.

Alternatively, NaHCO_3_ may mitigate fatigue by altering the intracellular and extracellular balance of strong ions such as potassium (K^+^), sodium (Na^+^), chloride (Cl^−^), and Ca^2+^ [26]. Indeed, NaHCO_3_ is suggested to maintain muscle excitability by increasing the influx of K^+^ into the muscle following ingestion and attenuating losses in intramuscular K^+^ during exercise [27–30]. A concomitant increase in muscular Cl^−^ uptake is also observed [27,29,30], which is suspected to work synergistically with K^+^ to protect force and excitability when muscles begin to depolarise [31]. This effect is further strengthened by an increased plasma Na^+^ [2,3,27,28], which together with changes in K^+^ and Cl^−^, could indicate an upregulation of Na^+^/K^+^-ATPase and Na^+^/K^+^/2Cl^−^-ATPase activity to limit depolarisation and preserve excitation-contraction coupling following NaHCO_3_ ingestion [16,30,31]. Currently, only Gough et al. have reported changes in the collective strong ion difference (SID) following NaHCO3 ingestion and whole-body exercise [24,27]. Interestingly, both studies significantly improved performance (4 km cycling time-trial: +1.4% [24], treadmill time-to-exhaustion: +28% [27]) with simultaneous increases in pre-exercise HCO_3_^−^ (+7.0 mmol·L^-1^ [24]; +8.8 mmol·L^-1^ [27]) and SID (+6 mEq·L^-1^ [24]; +10 mEq·L^-1^ [27]) when compared to placebo conditions. Given that the roles of H^+^ and acidosis are controversial in exercise performance [16–18], the resultant performance improvements could instead be attributed to the observed increases in the SID. However, neither of these studies purposely intended to identify a peak SID measurement, therefore it is currently unknown as to when the SID is maximised following NaHCO_3_ ingestion. It is plausible that if the SID peaks at a different time point in comparison to HCO_3_^−^ and pH, then a NaHCO_3_ ingestion strategy based upon a time to peak SID approach could optimise the SID mechanisms associated with performance enhancement.

Whether either of these mechanisms are ergogenic for young athletes remains unclear since adolescents have lesser reliance on glycolytic metabolism, lower amounts of muscle mass, and fewer type IIb muscle fibre recruitment than their adult peers [32–34]. These differences result in less H^+^ production at given exercise intensities, consequently suggesting that age-related differences may also exist in acid-base balance kinetics. Only Zajac et al. [35] have investigated the effects of NaHCO_3_ in adolescent athletes (age: 15.1 ± 0.4 yrs). The authors found that 0.3 g·kg BM^-1^ NaHCO_3_ improved 50 m swimming speed (+0.05 m·s^-1^, 2.6%) and reduced the time to complete a 4 x 50 m swimming protocol (^−^1.4 s, 1.3%) despite only a 3 mmol·L^-1^ increase in HCO_3_^−^ concentrations. This increase in HCO_3_^−^ is lower than typically observed in adults [2,3] and below the proposed +5 mmol·L^-1^ threshold for increased ergogenic potential [19,20]. However, as HCO_3_^−^ was measured at 60 min post-ingestion, and exercise commenced at 90 min post-ingestion, it is possible that further increases in HCO3^−^ occurred within this 30 min window [2]. Furthermore, HCO3^−^ was reported at the group level only, leaving it unknown as to whether some adolescents had greater elevations at this time point, or whether large inter-individual variations exist in the time to peak in accordance with adult-like NaHCO_3_ absorption [2,3,22]. The purpose of this study was therefore to explore the time course changes and peak blood concentrations of three acid-base balance variables (HCO_3_^−^, pH, and the SID) in highly trained adolescent athletes following the ingestion of 0.3 g·kg BM^-1^ NaHCO_3_.

## Materials and Methods

### Participants

Twelve national level swimmers from a performance swim programme volunteered to participate in this study (Table 1). Three of the swimmers had recently represented Great Britain at either the European Junior Swimming Championships or the European Youth Olympic Festival in 2019, whilst seven had qualified for the senior British Swimming Championships in April 2020. At the time of the study, all swimmers were ranked within the top 25 in Great Britain in at least one event within their respected age group (mean FINA points = 696 ± 62) and were completing a minimum of 7 x 2.5 hours pool-based and 2 x 1-hour land-based training sessions per week. The study was granted institutional ethical approval and both the swimmers and their parents/guardians provided written informed consent prior to their participation in the study.

**Table 1.** Anthropometric and performance characteristics of study participants (± SD)

| Characteristics | Males<br>(n = 5) | Females<br>(n = 7) | Combined<br>(n = 12) |
| --- | --- | --- | --- |
| Age (yrs) | 16.4 ± 1.1 | 15.6 ± 0.8 | 15.9 ± 1.0 |
| Height (m) | 1.78 ± 0.6 | 1.70 ± 0.4 | 1.73 ± 0.6 |
| Body mass (kg) | 72.4 ± 10.7 | 60.3 ± 4.8 | 65.3 ± 9.6 |
| Sum of 8 skinfolds (mm) | 55.2 ± 11.3 | 82.1 ± 21.5 | 70.9 ± 22.1 |
| Time as a competitive junior swimmer (yrs) | 8.2 ± 1.6 | 8.0 ± 0.8 | 8.1 ± 1.2 |
| **Personal Best Swimming Times (Freestyle – Long Course)** |  |  |  |
| 50 m (sec) | 26.0 ± 2.6 | 29.4 ± 1.5 | 28.0 ± 2.6 |
| 100 m (sec) | 54.7 ± 3.0 | 61.2 ± 1.4 | 58.8 ± 15.6 |
| 200 m (min : sec) | 1:57.7 ± 4.9 | 2:11.3 ± 3.6 | 2:05.6 ± 8.1 |

### Pre-Experimental Procedures

Each participant was required to attend the laboratory as per their usual training time (17:00 – 20:30 hrs) having eaten as they normally would prior to a competitive race (i.e., self-selected timing and meal composition). Previous studies have assessed time course changes in HCO_3_^−^ and pH in a fasted state [2,3,36] or following a standardised meal [22], however, these conditions do not replicate the variance in nutritional intakes that would take place in competition. This ingestion method was selected to consider the effects of individualised nutrition on baseline acid-base balance, electrolyte status, and absorption rate that would occur in practice [37]. In addition, the ingestion of a meal is also shown to reduce the occurrence and severity of potential gastrointestinal (GI) side-effects that are associated with NaHCO_3_ ingestion [38]. Similarly, water was permitted to be consumed *ad libitum* (mean intake: 1.2 ± 0.5 L, range: 0.5 – 2.0 L), though ingestion of further nutrients were restricted once NaHCO_3_ had been consumed and blood changes were being monitored. Participants were also asked to refrain from additional exercise outside of their regular swim training programme in the 48 hours prior to participating in the study. No swimmers had ingested caffeine, although the consumption of creatine (participants 5, 8, 9, 10, and 11) and beta-alanine (participants 9 and 10) were reported. Since co-ingestion of multiple ergogenic aids is common in high-level sport, these participants were included in subgroup analysis rather than excluded from the study altogether. All trials were carried out between December 2019 and February 2020 during a specific race preparation training period (weekly swimming volume = 50.9 ± 3.4 km).

### Protocol and Measurements

Upon arrival to the laboratory, participants engaged in five min of seated rest before a baseline capillary sample of whole blood was collected from the fingertip into a 100 μL sodium-heparin coated glass clinitube (Radiometer Medical Ltd., Copenhagen, Denmark). Blood samples were immediately analysed using a blood gas analyser (ABL9, Radiometer Medical Ltd., Copenhgen, Denmark) for measurements of HCO3^−^, pH, K^+^, Na^+^, Ca^2+^, and Cl^−^. An additional 5 μL sample was taken for the analysis of blood lactate (La^−^) (Lactate Pro 2, Arkray, Japan) which was used in the following formula to calculate the apparent SID using a freely available spreadsheet: K^+^ + Na^+^ + Ca^2+^ – Cl^−^ – La^−^ [39]. This method of calculating the apparent SID has been previously utilised in similar research investigating the SID alongside NaHCO_3_ ingestion [24,27]. Participants then ingested 0.3 g·kg^-1^ BM NaHCO_3_ (Dr. Oetker, Bielefeld, Germany) contained in gelatine capsules (1 g NaHCO_3_ per capsule; Size 00, Bulk Powders, Colchester, UK) within a five min period prior to 165 min of quiet seated rest. Blood samples were obtained and analysed on eight more occasions (60, 75, 90, 105, 120, 135, 150, and 165 min post-NaHCO_3_ ingestion) with further samples taken at 180 (n = 4) and 195 min (n = 1) to ensure a peak HCO_3_^−^ was found. No samples were taken within the first 60 min since 0.3 g·kg^-1^ BM NaHCO_3_ in capsule form was not expected to display peak HCO_3_^−^ concentrations in this time [2,22].

Gastrointestinal side-effects were monitored using nine 200 mm visual analogue scales (VAS) for nausea, flatulence, stomach cramping, belching, stomach ache, bowel urgency, diarrhoea, vomiting and stomach bloating consistent with previous NaHCO_3_ research [2,24,25,40]. The VAS scales were labelled “no symptom” on the left side and “severe symptom” on the right side.

## Statistical Analysis

Data was normally distributed (Shapiro-Wilks) and sphericity was assumed (Mauchly) prior to statistical analysis. If sphericity was violated, Huyn-Feldt (epsilon value >0.75) or Greenhouse-Geiser (epsilon value <0.75) corrections were applied. A repeated measures ANOVA was conducted to establish mean differences between the time to peak of the three alkalotic variables (HCO_3_^−^, pH, and the SID) and group mean differences for each blood metabolite (HCO_3_^−^, pH, SID, K^+^, Na^+^, Ca^2+^, and Cl^−^) at each blood sampling time point. Post hoc comparisons were determined by the Bonferroni correction and effect sizes are reported as partial eta squared (Pη^2^). A paired samples t-test was used to determine differences in the time to peak and absolute blood acid-base changes between the participants that co-ingested supplements (SUP) and those that had not been ingesting any other substances (NONE) during the investigation (n = 6 each group). Additional effect sizes were calculated where appropriate using Hedges’ *g* bias correction [41]. Effect size calculations were interpreted as trivial (*g* = <0.20), small (*g* = 0.20 – 0.49), medium (*g* = 0.50 – 0.79), and large (*g* = ≥0.80) [42]. Coefficient of variation (CV) was calculated and reported using SD/mean*100. All statistical tests were completed using Statistical Package for the Social Sciences (SPSS), version 25 (IBM, Chicago, USA). All data are reported as mean ± SD with statistical significance set at p <0.05.

## Results

There were no differences between the time to peak of each acid-base variable (HCO_3_^−^ = 130 ± 35 min, pH = 120 ± 38 min, SID = 96 ± 35 min; p = 0.063, Pη^2^ = 0.223). Whilst significance was not found, a large effect size was observed between the SID and HCO_3_^−^ (mean difference: 34 min, *g* = 0.91). A moderate effect size was also observed between time taken to reach a peak SID and pH (24 min, *g* = 0.63), whereas the effect size was small between HCO_3_^−^ vs. pH (10 min, *g* = 0.26). Large inter-individual variations were present in the time to peak (CV: HCO_3_^−^ = 27%; pH = 32%; SID = 37%; Fig 1) and absolute increase from baseline for each alkalotic measure (HCO_3_^−^ = +6.8 ± 2.4 mmol·L^-1^, CV = 35%; pH = +0.10 ± 0.04 units, CV = 42%; SID = +3.8 ± 2.3 mEq·L^-1^, CV = 61%; Table 2).

**Fig 1.**
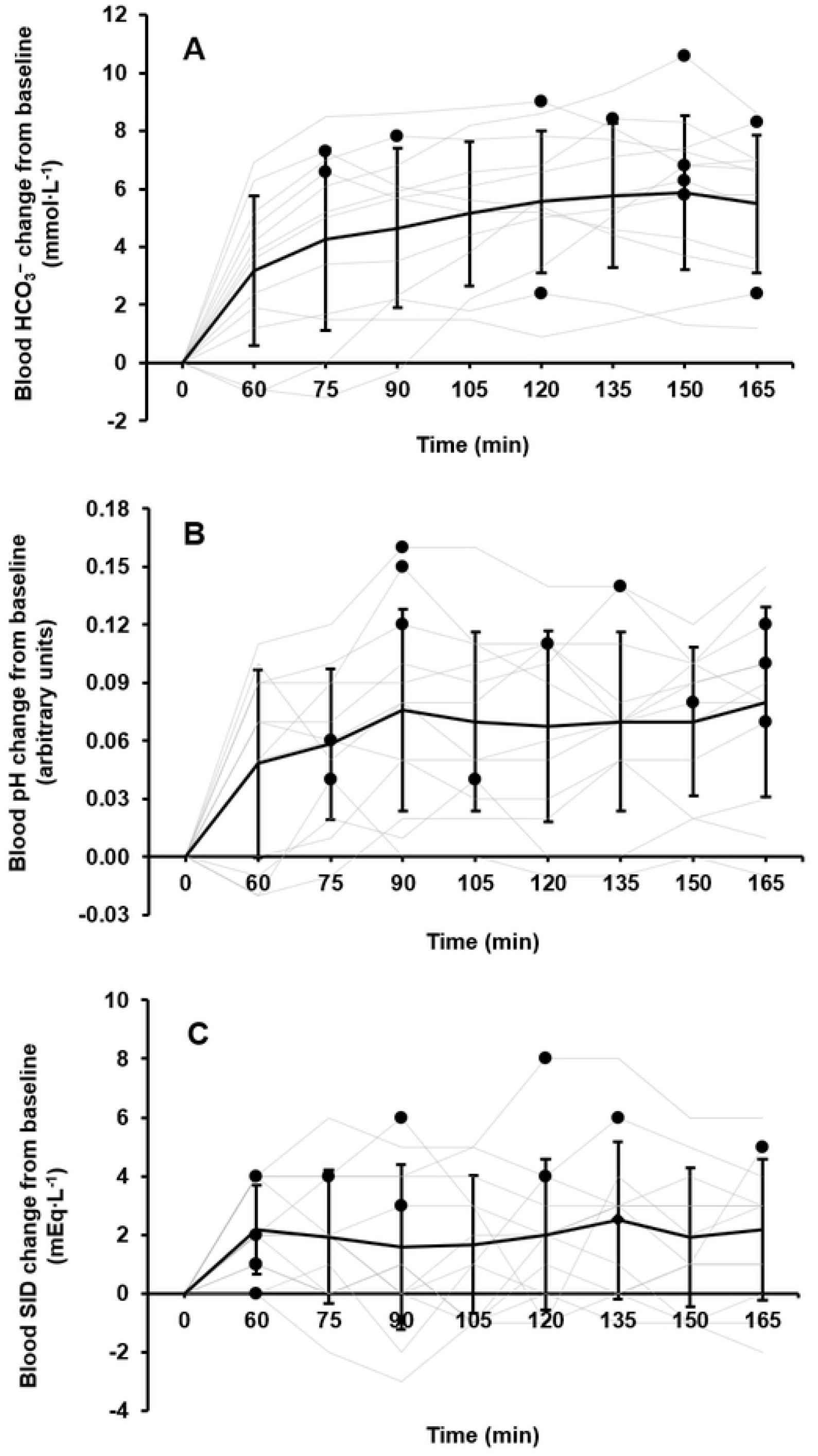
Individual dose-reponse (n = 5) for a) blood bicarbonate (HCO3^−^), b) pH, and c) the strong ion difference (SID) following 0.3 g·kg BM^-1^ of sodium bicarbonate (NaHCO_3_). Only five participants data is shown for clarity and to emphasise the different patterns that emerged in acid-base repsonses

**Table 2.** Individual metabolite responses to 0.3 g·kg BM^-1^ sodium bicarbonate (NaHCO_3_)

|  | Participant (Gender) |  |  |  |  |  |  |  |  |  |  |  |
| --- | --- | --- | --- | --- | --- | --- | --- | --- | --- | --- | --- | --- |
|  | 1 (M) | 2 (M) | 3 (F) | 4 (F) | 5 (M) | 6 (M) | 7 (F) | 8 (F) | 9 (F) | 10 (M) | 11 (F) | 12 (F) |
| **HCO<sub>3</sub><sup>-</sup> baseline (mmol·L<sup>-1</sup>)** | 26.5 | 28.0 | 23.3 | 23.9 | 26.0 | 25.6 | 24.8 | 25.9 | 24.7 | 26.4 | 26.4 | 23.6 |
| **HCO<sub>3</sub><sup>-</sup> peak (mmol·L<sup>-1</sup>)** | 34.3 | 35.3 | 32.3 | 34.5 | 32.8 | 33.9 | 31.4 | 28.3 | 30.5 | 32.7 | 34.8 | 26.4 |
| **Absolute change (mmol·L<sup>-1</sup>)** | +7.8 | +7.3 | +9.0 | +10.6 | +6.8 | +8.3 | +6.6 | +2.4 | +5.8 | +6.3 | +8.4 | +2.8 |
| **Time to peak (min)** | 90 | 75 | 120 | 150 | 150 | 165 | 75 | 120 | 150 | 150 | 135 | 180 |
| **pH baseline** | 7.41 | 7.46 | 7.40 | 7.40 | 7.40 | 7.41 | 7.39 | 7.41 | 7.41 | 7.40 | 7.38 | 7.43 |
| **pH peak** | 7.57 | 7.52 | 7.52 | 7.54 | 7.49 | 7.53 | 7.54 | 7.45 | 7.51 | 7.48 | 7.49 | 7.47 |
| **Absolute change** | +0.16 | +0.06 | +0.12 | +0.14 | +0.09 | +0.12 | +0.15 | +0.04 | +0.10 | +0.08 | +0.11 | +0.04 |
| **Time to peak (min)** | 90 | 75 | 90 | 135 | 180 | 165 | 90 | 105 | 165 | 150 | 120 | 75 |
| **SID baseline (mEq·L<sup>-1</sup>)** | 33 | 37 | 33 | 32 | 33 | 35 | 34 | 33 | 31 | 30 | 34 | 32 |
| **SID peak (mEq·L<sup>-1</sup>)** | 36 | 38 | 39 | 38 | 37 | 35 | 39 | 35 | 35 | 38 | 38 | 34 |
| **Absolute change (mEq·L<sup>-1</sup>)** | +3 | +1 | +6 | +6 | +4 | 0 | +5 | +2 | +4 | +8 | +4 | +1 |
| **Time to peak (min)** | 90 | 60 | 90 | 135 | 120 | 60 | 165 | 120 | 60 | 120 | 75 | 60 |

Across the sampling timeframe, marked increases in acid-base variables were observed (HCO3^−^: p = <0.001, Pη^2^ = 0.609; pH: p = <0.001, Pη^2^ = 0.529; SID: p = 0.030, Pη^2^ = 0.217; Fig 2). Blood bicarbonate was elevated 60 min post-ingestion (+3.2 ± 2.6 mmol·L^-1^, +13%, p = 0.046) and remained +4 – 6 mmol·L^-1^ until 165 min post-ingestion (all p < 0.030). At 105 min mean blood HCO_3_^−^ had increased by 5.2 ± 2.5 mmol·L^-1^ and remained elevated above the proposed +5 mmol·L^-1^ ergogenic threshold until the final 165 min time point (peak: 150 min, +5.9 ± 2.7 mmol·L^-1^, +23%). An increased pH occurred at 75 min post-ingestion (+0.06 ± 0.04 units, +0.8%, p = 0.010) and this level of increase was sustained at all remaining points in time (+0.06 – 0.08 units, all p < 0.030). Peak pH occurred at 165 min post-ingestion (+0.08 ± 0.05 units, +1.1%). The SID increased by 2.2 ± 1.5 mEq·L^-1^ (+7%, p = 0.017) at 60 min post-ingestion and peaked at 135 min post-ingestion (+2.5 ± 2.7 mEq·L^-1^, +8%), however, this did not reach statistical significance.

**Fig. 2.**
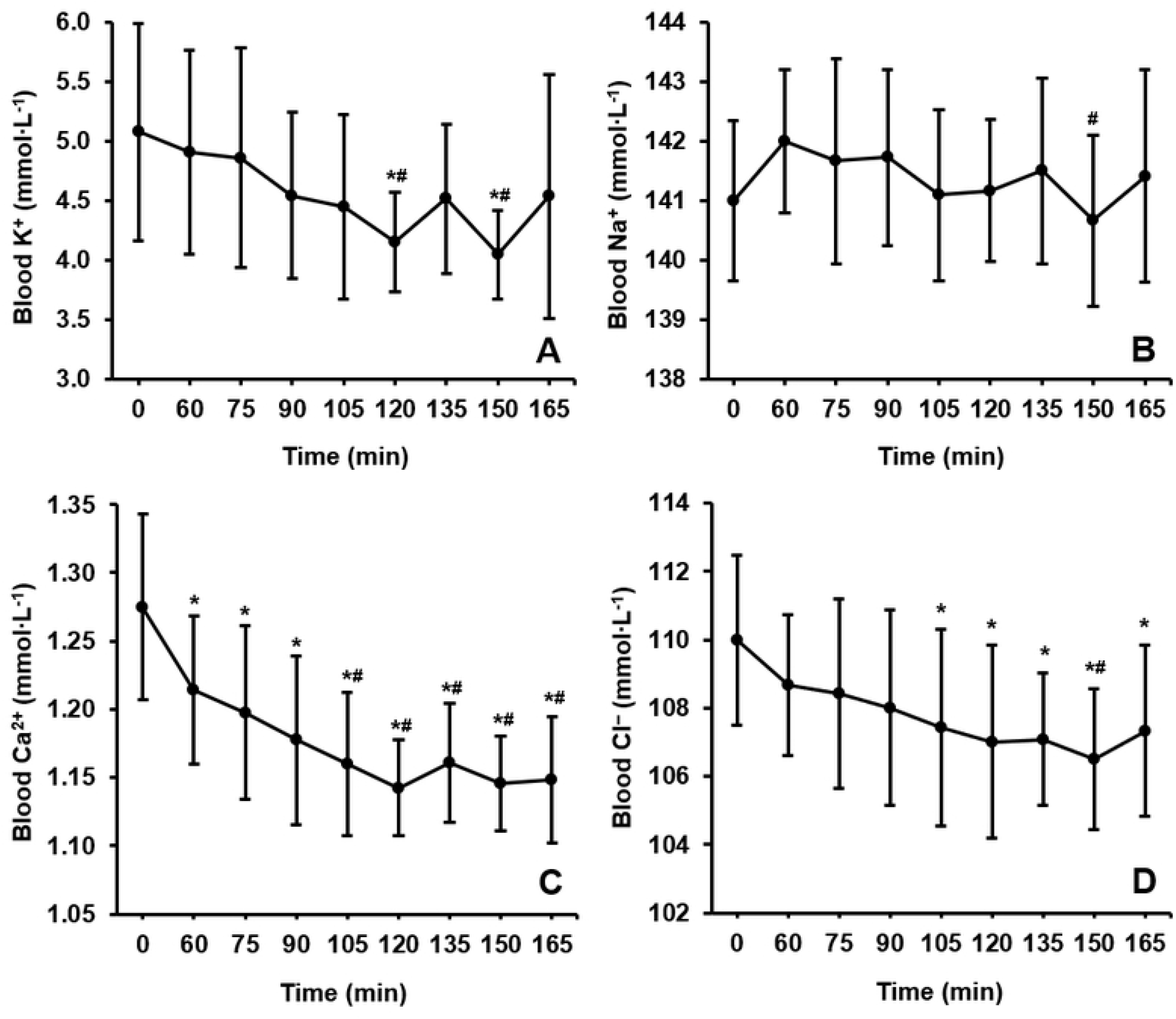
Mean dose-response changes in plasma a) blood bicarbonate (HCO3^−^), b) pH, and c) the strong ion difference (SID) following the ingestion of 0.3 g·kg of sodium bicarbonate (NaHCO3). * denotes a large effect size (*g* = >0.80) from baseline. # denotes a large effect size (*g* = >0.80) compared to 60 min post-ingestion

Ingestion of NaHCO_3_ produced ionic shifts in the balance of strong ions (K^+^: p = 0.005, Pη^2^ = 0.215; Na^+^: p = 0.044, Pη^2^ = 0.160; Ca^2+^: p = <0.001, Pη^2^ = 0.562; Cl^−^: p = 0.016, Pη^2^ = 0.306). The electrolytes with the greatest absolute changes were K^+^ and Cl^−^. Large inter-individual variations restricted the ability to detect statistical significances for all metabolites, therefore group level results are described using effect sizes (Fig 3). This analysis showed that K^+^ reduced gradually from baseline before reaching peak declines at 120 (–0.93 ± 1.07 mmol·L^-1^, –18%, *g* = 1.26) and 150 min (–1.03 ± 0.93 mmol·L^-1^, –20%, *g* = 1.64) post-ingestion. Large effect sizes were calculated for the reductions in Cl^−^ from 105 to 165 min post-ingestion (all *g* > 0.80) with the lowest point also occurring at 150 min post-ingestion (–3.5 ± 1.9 mmol·L^-1^, –3%, p = 0.002). The point in time in which each individual participant reached their lowest values of K^+^ (133 ± 28 min, CV = 21%), Cl^−^ (124 ± 27 min, CV = 21%), and Ca^2+^ (141 ± 32 min, CV = 23%) averaged within 20 min of one another, albeit with large inter-individual variances. Conversely, Na^+^ was largely unchanged throughout the sampling period. The absolute changes from baseline also displayed high inter-individual variation for each variable (K^+^: –1.28 ± 0.88 mmol·L^-1^, CV = 69%; Cl^−^: – 4.5 ± 2.1 mmol·L^-1^, CV = 47%; Ca^2+^: ^−^0.16 ± 0.07 mmol·L^-1^, CV = 43%; Na^+^: +1.6 ± 1.3 mmol·L^-1^, CV = 83%).

**Fig 3.** Change in mean plasma concentrations of a) potassium (K^+^), b) sodium (Na^+^), c) calcium (Ca^2+^), and d) chloride (Cl^−^) following the ingestion of 0.3 g·kg BM^-1^ of sodium bicarbonate (NaHCO_3_). * denotes a large effect size (*g* = >0.80) from baseline. # denotes a large effect size (*g* = >0.80) compared to 60 min post-ingestion

Participants that were consuming creatine and beta-alanine (SUP, n = 6) at the time of the study were younger than those intaking no other (NONE, n = 6) supplements (15.2 ± 0.4 vs. 16.7 ± 0.8 yrs, p = 0.001), however, both groups ingested a similar total amount of NaHCO_3_ (19.7 ± 1.9 vs. 20.1 ± 3.8 capsules, p = 0.474). Absolute increases from baseline were different for HCO_3_^−^ (SUP: +5.4 ± 2.4 mmol·L^-1^, NONE: +8.3 ± 1.4 mmol·L^-1^, p = 0.017, *g* = 1.43) and pH (SUP: +0.08 ± 0.03 units, NONE: +0.13 ± 0.04 units, p = 0.006, *g* = 1.37), but not for the SID (SUP: +4.0 ± 2.2 mEq·L^-1^, NONE: +3.5 ± 2.6 mEq·L^-1^, p = 0.656, *g* = 0.20). No difference was found in the time of peak measurements in pH (SUP: 133 ± 40 min, NONE: 108 ± 35 min, p = 0.289, *g* = 0.64) or the SID (SUP: 93 ± 31 min, NONE: 100 ± 42 min, p = 0.762, *g* = 0.18). Similarly, statistical significance was not found for the time that peak HCO_3_^−^ occurred (SUP: 148 ± 20 min, NONE: 113 ± 39 min, p = 0.128), although a large effect size was calculated between peak time points (*g* = 1.09). Baseline concentrations of the SID were higher in the non-supplement group (34.0 ± 1.8 vs. 32.2 ± 1.5 mEq·L^-1^, p = 0.020), but there were no differences for HCO3^−^ (SUP: 25.5 ± 1.1 mmol·L^-1^, NONE: 25.4 ± 1.7 mmol·L^-1^, p = 0.839) or pH (SUP: 7.41 ± 0.02 units, NONE: 7.41 ± 0.02 units, p = 0.625).

All participants reported at least one GI side-effect, though the severity of these instances was rated ≤5/10 in 94% (29/31) of occurrences (Table 3). Of the reported side-effects, 71% (22/31) of instances peaked in severity at 60 and 75 min post-ingestion. The most common symptom was belching (58%, 7/12), followed by stomach bloating and nausea (50%, 6/12).

**Table 3.** The three most severe symptoms of gastrointestinal discomfort experienced following the ingestion of 0.3 g·kg BM^-1^ sodium bicarbonate (NaHCO_3_)

| Participant<br>(Body mass) | Caps | Symptom 1 | Severity<br>(n/10) | Peak<br>(mins) | Symptom 2 | Severity<br>(n/10) | Peak<br>(mins) | Symptom 3 | Severity<br>(n/10) | Peak<br>(mins) | Aggregated |
| --- | --- | --- | --- | --- | --- | --- | --- | --- | --- | --- | --- |
| 1 (90.7 kg) | 28 | Belching | 4.9 | 60 | Stomach<br>bloating | 4.9 | 60 | Nausea | 2.3 | 60 | 17.7 |
| 2 (69.5 kg) | 21 | Belching | 2.7 | 60 | Stomach<br>bloating | 1.7 | 135 | Stomach<br>cramp | 1.1 | 60 | 6.6 |
| 3 (55.8 kg) | 17 | Nausea | 4.1 | 75 | Stomach<br>bloating | 1.4 | 60 | None | 0.0 | N/A | 5.5 |
| 4 (62.2 kg) | 19 | Belching | 3.9 | 60 | Stomach<br>bloating | 2.4 | 60 | Nausea | 0.7 | 75 | 6.9 |
| 5 (72.1 kg) | 22 | Belching | 1.7 | 75 | None | 0.0 | N/A | None | 0.0 | N/A | 1.7 |
| 6 (64.6 kg) | 20 | Stomach<br>ache | 4.0 | 165 | None | 0.0 | N/A | None | 0.0 | N/A | 4.0 |
| 7 (60.6 kg) | 19 | Nausea | 1.4 | 60 | None | 0.0 | N/A | None | 0.0 | N/A | 1.4 |
| 8 (54.3 kg) | 17 | Belching | 4.9 | 60 | Flatulence | 3.9 | 165 | Stomach<br>ache | 1.5 | 135 | 11.0 |
| 9 (57.7 kg) | 18 | Nausea | 2.6 | 60 | None | 0.0 | N/A | None | 0.0 | N/A | 2.6 |
| 10 (65.0 kg) | 20 | Belching | 5.0 | 60 | None | 0.0 | N/A | None | 0.0 | N/A | 5.0 |
| 11 (67.9 kg) | 21 | Flatulence | 2.6 | 150 | Stomach<br>bloating | 2.1 | 60 | Bowel<br>urgency | 1.5 | 165 | 6.1 |
| 12 (63.7 kg) | 20 | Vomiting | 10.0 | 105 | Nausea | 10.0 | 105 | Stomach<br>ache | 1.5 | 90 | 22.3 |

261 Aggregated = Sum of the most severe incidence of all nine symptoms (n/90). Caps = Number of capsules consumed (1 g of NaHCO<sub>3</sub> per capsule)

## Discussion

This was the first study to compare the time to peak of three methods of measuring peak alkalosis (i.e., HCO_3_^−^, pH, and the SID) following the ingestion of 0.3 g·kg BM^-1^ NaHCO_3_ in highly trained adolescent male and female swimmers. Though no statistical significance was observed between the three different measures, there was a difference of 34 min separating the time in which HCO_3_^−^ and the SID peaked within the blood. In addition, large inter-individual variation was observed in the time to peak of each acid-base balance variable, whereas the co-ingestion of other nutritional ergogenic aids blunted both the absolute increase and time to peak of blood HCO_3_^−^. Erratic time course behaviour in HCO_3_^−^ was also observed, whereby some individuals sustained increases >5 mmol·L^-1^ throughout the 165 min sampling period, while others had a rapid decline once a peak concentration occurred. Collectively, these findings contrast previous suggestions of a long lasting ‘ergogenic window’ following the ingestion of NaHCO_3_ capsules, and therefore reiterate the necessity of an individual time to peak approach in applied practice when conditions are not controlled (e.g., food, fluid, and supplement intakes). Future research should aim to clarify the impact of individualised ingestion strategies, including a time to peak SID approach, on exercise performance when athletes are following their real-world nutrition practices in order to elucidate the optimal use of NaHCO_3_ in competition.

On an individual basis, peak HCO_3_^−^ and pH concentrations occurred within 30 min of one another for 11 of 12 participants. The close relationship between these two measurements is synonymous with previous research exploring the dose-response relationship of NaHCO_3_ consumed in gelatine capsules [3,22,38]. The remainder of this discussion will therefore focus on HCO_3_^−^ as this measurement typically displays less variance and a greater degree of reliability when compared with pH [2,21]. The time taken for trained adolescents to reach peak HCO_3_^−^ concentrations ranged considerably within the present study (75 – 180 min), which is in accordance with similar research (40 – 240 min) when recreational adults ingested NaHCO_3_ capsules in fed or fasted states [3,21,22]. The reason for the inter-individual variation remains elusive, though it is postulated that genetic differences in the rate of gastric emptying, intestinal motility, and gastrointestinal blood flow may contribute by influencing absorption characteristics [43,44]. Another factor that appears to affect HCO_3_^−^ kinetics is the co-ingestion of buffering supplements (i.e., creatine and beta-alanine), as these agents appeared to delay the time to peak by approximately 35 min whilst concomitantly mitigating the absolute increases in HCO_3_^−^ (+5 vs. +8 mmol·L^-1^) compared to the adolescents ingesting no other supplements. While this finding is confounded by a small sample size (n = 6) and a greater age and potentially maturational status (+2 yrs) of the non-supplement subgroup, it is important to consider that highly trained adolescents will likely be consuming multiple ergogenic aids at times of competition [45]. Sports nutritionists should therefore aim to seek the time to peak of their athletes whilst they are in full competition preparation to minimise variations at competitive events.

The SID of highly trained adolescent swimmers was increased by NaHCO_3_ ingestion, albeit with a large individual variability in the time to peak (60 – 165 min). This finding supports research by Gough et al. [24,27] whereby an elevation in the SID was detected in accordance with performance enhancement in trained adults. Though it is somewhat surprising that Gough et al. reported greater group mean increases in the SID considering that neither of these studies purposely sought to identify a peak concentration (+6 and +10 vs. +4 mEq·L^-1^). Baseline (33 vs. 36 and 38 mEq·L^-1^) and absolute peak SID (37 vs. 42 and 46 mEq·L^-1^) was also lower in the present study, possibly inferring that an age-related difference exists in strong ion movements. Indeed, the current results somewhat support this notion since baseline SID was higher in the older co-supplementing subgroup (+2 mEq·L^-1^) despite only a small increase in age (16.7 ± 0.8 vs. 15.2 ± 0.4 yrs). More controlled settings are required to confirm this speculation, however, since a pre-ingestion meal and *ad libitum* water consumption could have affected baseline electrolyte balance [46], NaHCO3 absorption characteristics [47], and plasma volume [48] between each participant. For example, the observed increases in plasma Na^+^ appeared to be mitigated (+1.6 vs. +3.0 mmol·L^-1^) in comparison with previous research investigating NaHCO3 consumption in a fasted state [2,3,27]. Nonetheless, large effect sizes were calculated for plasma reductions of K^+^, Ca^2+^, and Cl^−^ between 120 – 150 min post-ingestion that could infer the intramuscular uptake of these ions [49]. However, since this study did not consider changes in the intramuscular SID or plasma volume, future research is required to elucidate the role of NaHCO_3_ on the SID.

Consuming an individualised meal at a self-selected time prior to the ingestion of NaHCO_3_ capsules successfully diminished the severity of GI discomfort within adolescent athletes. Whilst each participant recorded at least one side-effect, these were considered minor in 94% (29/31) of incidences. This supports research by Carr et al. [38] which found that the ingestion of NaHCO_3_ capsules alongside a 1.5 g·kg BM^-1^ carbohydrate meal lessened GI disturbances compared to other ingestion methods. This is suggested to slow the rate of NaHCO_3_ reduction to Na^+^ and carbonic acid (H_2_CO_3_), thereby lessening the rate of acid-base alterations that occur within the stomach (e.g., increases in carbon dioxide and water) [50,51]. The lack of severe GI disturbances could also be related to the low body mass of the participant cohort, and therefore reduced absolute amount of total NaHCO_3_ ingested. For instance, the current study and Zajac et al. [35] found limited GI side-effects in adolescent swimmers weighing 65 ± 10 kg and 56 ± 1 kg, respectively, whereas diarrhoea and/or vomiting occurred in over 50% of senior rugby players with a body mass of 95 ± 13 kg [40]. Consequently, it may be that the current ingestion method may only alleviate GI disturbances in athletes of body mass (i.e., <70 kg).

The study methods were applied based on the logistical (e.g., taking highly trained swimmers out of training) and ethical considerations of the participant cohort (e.g., repeated blood analysis of young athletes). This limited the current study from including a placebo experiment or repeatability measures which could have substantially altered the interpretation of results. Previous work analysing blood analyte time course changes using capillary bloods detected no changes from baseline for either HCO3^−^, pH, or Na^+^ when consuming a placebo treatment [2], whereas the time in which HCO3^−^ peaks in the bloodstream was considered to have a good repeatability (technical error = 14 min, ICC = 0.77) in highly trained athletes consuming 0.3 g·kg BM^-1^ NaHCO_3_ in capsules [21]. However, it should be noted that both of these studies analysed blood variables under fasted conditions. Alternatively, de Oliveira et al. [22] identified that the repeatability of the peak HCO_3_^−^ timing was poor (technical error = 39 min, ICC = 0.34) when 0.3 g·kg BM^-1^ NaHCO_3_ capsules were consumed with a standardised meal. This stark difference was likely the outcome of methodologies used (e.g., venous vs. capillary samples, 240 vs. 180 min sampling period), therefore it is unclear to what extent the inclusion of an uncontrolled pre-ingestion meal had upon the repeatability measures in the present study. Nonetheless, the main purpose of this research was to identify whether any practical differences occur between three different measures of blood acid-base balance when highly trained adolescent athletes consumed NaHCO_3_ supplementation. As a result, this is the first study to identify an individual time to peak SID, and future research should investigate whether ingesting NaHCO_3_ at this alternate time point could be a more ergogenic for exercise performance versus other contemporary ingestion strategies (e.g., time to peak HCO_3_^−^, fixed ingestion timing).

The process of identifying a true peak HCO_3_^−^ measurement has recently been criticised when using NaHCO_3_ capsules since these produce large increases in HCO_3_^−^ (>5 mmol·L^-1^) for periods exceeding 100 min in duration [22,23]. This suggests that a proposed ‘ergogenic window’ could mitigate the necessity of an isolated time point in practice. Results from the present study contrast this assumption, however, since less 50% of participants had an increase in HCO_3_^−^ in excess of 5 mmol.L^-1^ which lasted for more than 90 min. Moreover, this window occurred at a completely separate point in time for some participants (i.e., 60 – 120 min, 135 – 165 min post-ingestion), whereas others had erratic time course changes which included decreases in HCO_3_^−^ concentration recorded at 75 – 90 min post-ingestion. Due to the findings of the current study, it is still recommended that the individual time point of peak alkalosis is required within a highly trained adolescent cohort. Furthermore, these data also support the notion that the traditional, fixed time point method of ingestion is flawed within adolescents, given that large inter-individual variations were apparent within all three methods of determining peak alkalosis. It is therefore suggested that adolescent athletes undergo testing to determine time to peak alkalosis prior using NaHCO_3_ supplementation at competitive events. However, considering that this is an expensive procedure, adolescents should remain cautious since the effects of individualised ingestion methods on exercise performance are currently untested within this cohort.

## Conclusion

This study shows that highly trained adolescent athletes have marked increases and highly individual time course changes in blood HCO_3_^−^, pH, and SID following the ingestion of NaHCO_3_ capsules. Importantly, the post-ingestion time point in which the individual time to peak occurred for HCO_3_^−^ and the SID was separated by a large effect size. Given that the effects of acidosis on exercise performance are controversial, this finding suggests that using a time to peak SID approach could be a more appropriate NaHCO_3_ ingestion strategy in practice. Consequently, future research should directly compare ingestion strategies based upon individual time to peak HCO_3_^−^ and SID to elucidate whether either approach has an any further ergogenic benefits on exercise performance. In addition, this study refutes the claim that NaHCO_3_ capsule ingestion produces a long lasting ‘ergogenic window’, supporting the need for adolescent athletes to identify their individual time to peak prior to use. It is acknowledged that a lack of control surrounding pre-ingestion food, fluid, and supplement intakes likely influenced the findings of this study, however, these conditions better replicate the actual changes that would occur when athletes are in full competition preparation.

## Acknowledgements

We gratefully acknowledge all participants who volunteered for the study and the City of Birmingham Swimming Club for facilitating the research process.

